# Monomethyl branched-chain fatty acid mediates amino acid sensing by mTORC1

**DOI:** 10.1101/2020.11.30.403246

**Authors:** Mengnan Zhu, Fukang Teng, Na Li, Li Zhang, Jing Shao, Haipeng Sun, Huanhu Zhu

## Abstract

Animals have developed various nutrient-sensing mechanisms for survival under fluctuating environments. Although extensive cultured cell-based analyses have discovered diverse mediators of amino acid sensing by mTOR, studies using animal models to illustrate intestine-initiated amino acid sensing mechanisms under specific physiological conditions are lacking. Here we developed a *Caenorhabditis elegans* model to examine the impact of amino acid deficiency on development. We discovered a leucine-derived monomethyl branched-chain fatty acid, and downstream glycosphingolipid, that critically mediate overall amino acid sensing by intestinal and neuronal mTORC1 that, in turn, regulates postembryonic development partly by controlling protein translation and ribosomal biogenesis. Additional data suggest that a similar mechanism may be conserved in mammals. This study uncovers an unexpected amino acid sensing mechanism mediated by a lipid biosynthesis pathway.

## Main Text

The ability to sense amino acid (AA) deficiency and make adjustments in cellular and developmental programs is fundamental for animals to survive under unfavorable environmental conditions^1,2^ Extensive past research has ascribed mTORC1 (mechanistic Target of Rapamycin Complex 1) as the major signaling system to sense AA. Mechanistic studies in the last decade, largely based on analyses in cultured cells, have indicated diverse mechanisms involving specific AA binding proteins and regulators that mediate AA sensing by mTORC1^3–7^ However, due to limited studies using animal models, how mTORC1 senses AA deficiency to regulate animal growth and development under specific physiological conditions remains largely unclear^8^. The complexity of generating AA deprivation conditions in live animals and the network of downstream activities in different tissues are among issues that render studies using animal models both more challenging and informative.

The nematode *Caenorhabditis elegans (C. elegans)* is an excellent animal model to study the regulation of various physiological functions, including nutrient sensing, primarily due to the genetic tools that can be applied. The combination of genetic manipulation (both in *C. elegans* and bacteria), food deprivation, and nutrient supplementation have permitted scientists to effectively reduce the level of certain nutrients and uncover various signaling mechanisms that regulate development and behaviors in response to the nutrient level change^9–17^ However, studies of AA sensing in *C. elegans* have been rather limited^18^ because of the technical difficulty in depriving AA from the laboratory food source (live *E. coli)* and developing assays that distinguish the roles of AA as a signal from their essential role in protein synthesis.

To address this problem, we designed an new nutrient-sensing assay in *C. elegans*, which was inspired by the standard AA sensing assay in cultured cells^8^. We created a dietary restriction condition by first diluting the common laboratory diet of *C. elegans (E. coli* OP50) and feeding it to the worms on antibiotic-treated NGM plates (to inhibit bacterial growth). This preparation had been previously used as a dietary restriction condition (previously called solid DR, we simply call it DR hereafter) for adult *C. elegans* to observe DR-induced lifespan extension^19^. When this DR condition was applied to newly hatched *C. elegans,* we found that the larvae grew slowly and most arrested at the L3 stage (Fig. 1a-d). Most animals survived and restored their normal development to gravid adults when transferred back to standard lab diet (considered as a favorable nutrient condition) (Fig. 1c, d, see Method). This reversible diapause suggests that such DR food presents a poor-quality food signal for developing larvae that *C. elegans* can sense and temporarily halt development as a protective measure.

**Fig 1.**
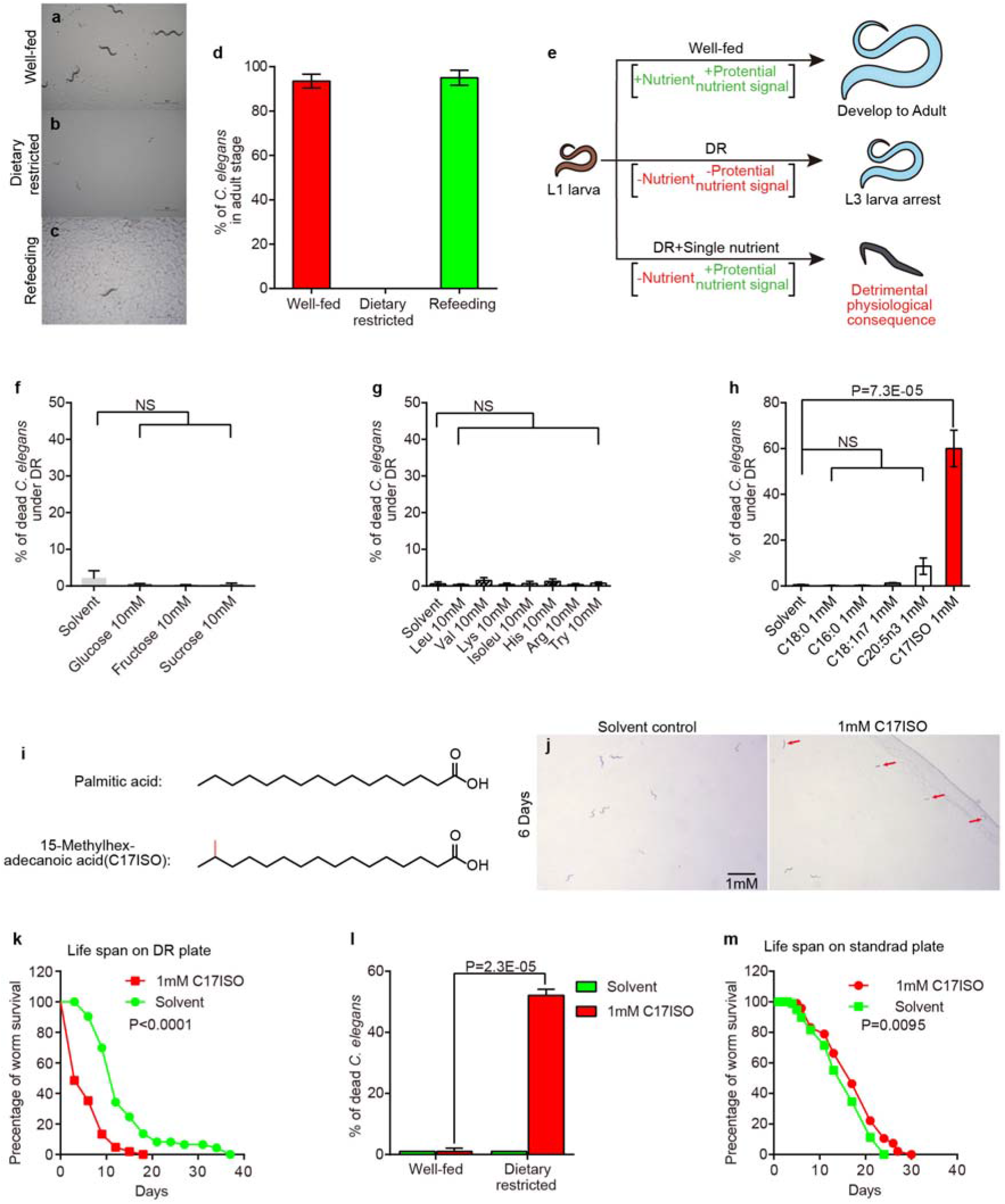
Dietary mmBCFAs triggered abnormal lethality under the nutrient restricted condition. **a-c**, Microscope images showing that WT *C. elegans* under the dietary restriction (DR) condition arrested growth at L3, whereas WT *C. elegans* under the favorable nutrient condition or switched from DR to the favorable nutrient condition grew to gravid adults. **d**, Percentage of animals at different developmental stages under the two conditions. **e**, Cartoon illustration showing the screen strategy to find key nutrient in the *C. elegans* nutrient-sensing process under the DR condition. **f-h**, Percentage of death in *C. elegans* supplied with indicated nutrients under the DR condition. Single essential amino acids (standard abbreviation) were used in (**g**). **i**, Chemical structures of palmitic acid and C17ISO. **j**, Dissection scope photos showed C. elegans grow under DR condition with or without C17ISO supplement. Arrowheads marked the dead animals. **k**, Survival curves showing C17ISO supplement reduced the survival rate of WT *C. elegans* grew under the DR condition. **l**, Bar graphs showing that percentage of death in *C. elegans* is dramatically increased with the supplementation of C17ISO under the DR (not under favorable nutrient condition. **m**, Survival curves showing the lifespan of *C. elegans* WT adults is not significantly changed with the C17ISO supplement under the favorable nutrient condition (standard plates). p=0.0095.

We then asked if adding a single dietary nutrient (carbohydrates, amino acids and fatty acids) could overcome this protective, DR-induced larval arrest. We reasoned that if the nutrient response system would rely on sensing one particular nutrient under the DR condition, then addition of such a nutrient might trigger further developmental activity with detrimental consequences (lethality) due to insufficiency of other nutrients (Fig. 1e). We first screened several simple saccharides (glucose, fructose and sucrose) and amino acids and found none caused a significant increase in lethality under DR (Fig. 1f,g). These data indicate that although these classical macronutrients (such as leucine or arginine) were reported to be able to activate the nutrient-sensing pathway under unfavorable nutrient conditions in the cell culture^5–7^, individually they are not sufficient to trigger a similar process in developing *C. elegans* under the DR condition.

We then screened several endogenously enriched fatty acids (including stearic acid[C18:0], palmitic acid [C16:0], cis-vaccenic acid [C18:1n7], eicosapentaenoic acid [C20:5n3], and 15-methylhexadecanoic acid [C17ISO])^20,21^ (Fig. 1h). We were surprised to find that only supplementation with C17ISO (Fig. 1i), a major monomethyl branched-chain fatty (mmBCFA) in *C. elegans*^9,22^, caused severe lethality under DR (Fig. 1h). About 60% of C17ISO supplemented animals were dead at the L3 stage in six days under the DR condition (Fig. 1h, j)(Method). The remaining animals were also very unhealthy (Fig. 1j). To confirm this finding, we measured the lifespan of C17ISO fed animals and found it was much shorter than the control group under DR (Fig. 1k). These data suggest that a single fatty acid, C17ISO, dramatically disrupted the protective arrest under the DR condition. We abbreviate such abnormal lethal phenotype as C17ISO-induced DUPED (Death Under Poor Environmental Diet).

mmBCFAs are essential for animal development and no adverse phenotypes caused by mmBCFA supplementation have been reported^9,22–27^. Therefore, it is unlikely that C17ISO-induced DUPED was caused by toxicity. To confirm this, we supplied well-fed *C. elegans* (grown on concentrated OP50 *E. coli* lawn, see method) with 1mM C17ISO and did not observe the DUPED phenotype (Fig. 1l). In addition, C17ISO supplementation also did not shorten the lifespan of well-fed *C. elegans* (Fig. 1m). These data further support that dietary C17ISO itself is not generally toxic and it causes lethality specifically under the DR condition. We thus hypothesized that C17ISO may act as a “master signal of nutrient” that overrides the protective arrest under the DR condition. Our further analyses showed that late larval stage and young adult *C. elegans* were most vulnerable to C17ISO-induced DUPED, while older adults were relatively resistant to the condition even though C17ISO still significantly shortened their lifespan (Extended Data Fig. 1a-c). The phenotypic difference could be due to a higher demand for nutrients during development. Thus, these data indicate that *C. elegans* critically evaluates a highly demanded nutrient by the level of mmBCFA during development, and the animal is able to sense the insufficiency of this nutrient and enter a protective arrest under DR.

We then reasoned that the shortage of this nutrient is the direct cause of C17ISO-induced DUPED and adding back the missing nutrient would suppress the lethality. We first excluded nucleotides, a highly demanded nutrient specifically in developing germlines, as a possible candidate because we found suppressing or enhancing the proliferation of *C. elegans* germ cells by Notch receptor *glp-1* loss-of-function and gain-of-function mutants (which critically mediates the nucleotide sensing and germline expansion^28^) did not affect C17ISO-induced DUPED (Extended Data Fig. 2a,b). Next we tested potential sources of energy, glucose or palmitate (C16:0), but they did not suppress the lethality (Fig. 2a). In contrast, supplementation with an AA mixture almost completely suppressed the C17ISO-induced DUPED lethality and restored the growth of about 40% of arrested animals to adulthood (Fig. 2b,c). These data suggest that amino acid deficiency may be the main cause of C17ISO-induced DUPED, and that dietary supplementation of C17ISO, but no single essential AA (Fig. 1g), was sensed by *C. elegans* as the signal for sufficient amino acids under DR. In other words, C17ISO may mediate overall AA sensing in *C. elegans* under the DR condition. In addition, supplementation of either the mixture of essential amino acids (EAA) (EAA: leucine, isoleucine, valine, phenylalanine, arginine, histidine, lysine, tryptophan, methionine and threonine)^29^, or mixture of non-essential amino acids (NEAA: consists of the rest of nine amino acids except asparagine) could not restore the development (Fig. 2C). Interestingly, we found that supplementation with the NEAA mix completely suppressed C17ISO-induced DUPED, while supplementation with the EAA mix significantly enhanced the DUPED phenotype (Fig. 2d). These data suggest that C17ISO-induced DUPED is mainly due to insufficient NEAA in our DR condition. The enhancement of the DUPED phenotype by supplementation with EAA might be due to an upregulation of the biosynthesis of endogenous mmBCFA (which is biosynthesized from the EAA leucine) or its downstream metabolite. Taken together, these data indicate that mmBCFA C17ISO (either from the diet or endogenous biosynthesis) acts as the key mediator of the overall amino acid level and signals to a nutrient response system to regulate the larval development program in *C. elegans* larvae.

**Fig 2.**
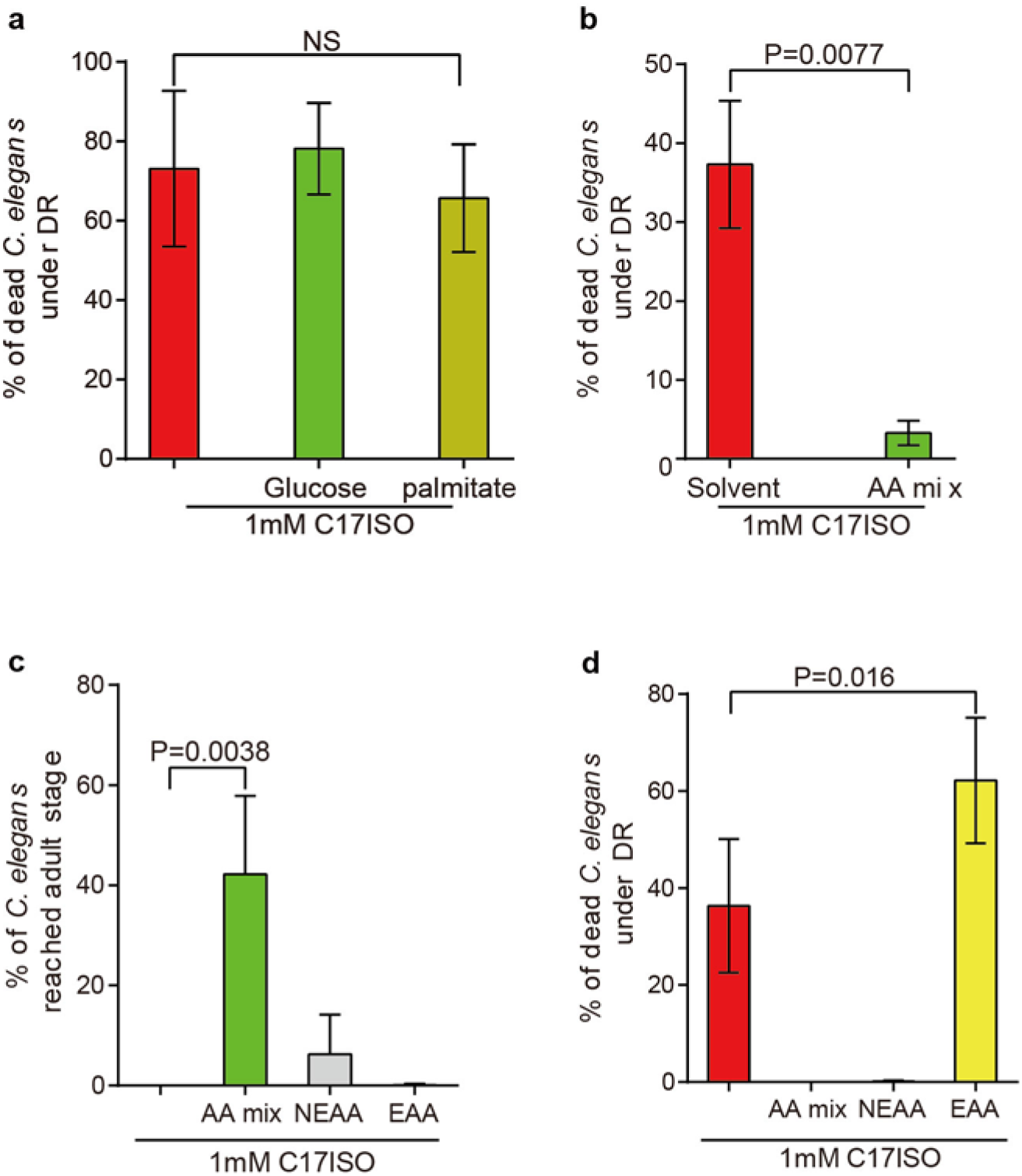
Amino acid shortage is the major cause of C17ISO-induced lethality under the DR condition. **a**, Bar graphs showing that percentage of death in *C. elegans* induced by C17ISO supplementation under the DR condition (C17ISO-induced DUPED) was not affected by the addition of glucose or palmitate. **b**, Bar graphs showing that C17ISO-induced DUPED is almost fully suppressed by the addition of the mixture of AA (AA mix). **c**, Bar graphs showing the impact of AA supplementation on the percentage of *C. elegans* reached adulthood under the DR and C17ISO supplementation condition. Recovery of animal growth was observed only with the supplementation of AA mix, but not with the mixture of essential (EAA) or nonessential amino acids (NEAA). **d**, Bar graphs showing that C17ISO-induced DUPED (red bar) was fully suppressed by the addition of NEAA Mix but enhanced by the addition of EAA mix.

Next, we investigated the mechanism by which C17ISO causes the DUPED phenotype under DR. We first turned to the mmBCFA synthesis pathway. In *C. elegans,* dietary C17ISO was previously reported to be metabolized to 13-Methyltetradecanoic acid (C15ISO) that is used to synthesize sphingolipids (Fig. 3a)^26,27,30^, which are essential for promoting early postembryonic development and foraging behaviors under the favorable nutrient condition^23,26^. We first determined that dietary C15ISO also induced the DUPED phenotype (Fig. 3b). Moreover, blocking the alpha hydrolase of very long chain fatty acids by *fath-1* RNAi, or ceramide glucosyltransferases by *cgt-1/2/3(-);nprl-3(-)[we* used the *nprl-3(-)* to bypass *cgt-1/2/3(-)* caused L1 arrest, see Method] also showed pronounced suppression of C17ISO-induced DUPED (Fig. 3a,c,d). However, blocking mannosylglucosylceramide transferase by *bre-3(-),* the next modification of glucosylceramide, did not cause suppression (Fig. 3a,e). These data indicate that C17ISO-derived d17isoGlcCer, in addition to its role in early development, also plays a major role in mediating C17ISO-induced DUPED under the DR condition. Interestingly, an isoleucine-derived mmBCFA 14-methylhexadecanoic acid (C17anteISO), which has high structural similarity to C17ISO and is able to fully suppress mmBCFA deficiency-caused early postembryonic developmental arrest^22,26^, could not trigger the DUPED phenotype at all (Extended Data Fig. 3a). These data indicate that C17ISO/d17isoGlcCer induced DUPED via a different mechanism from its role in promoting the early postembryonic development.

**Fig 3.**
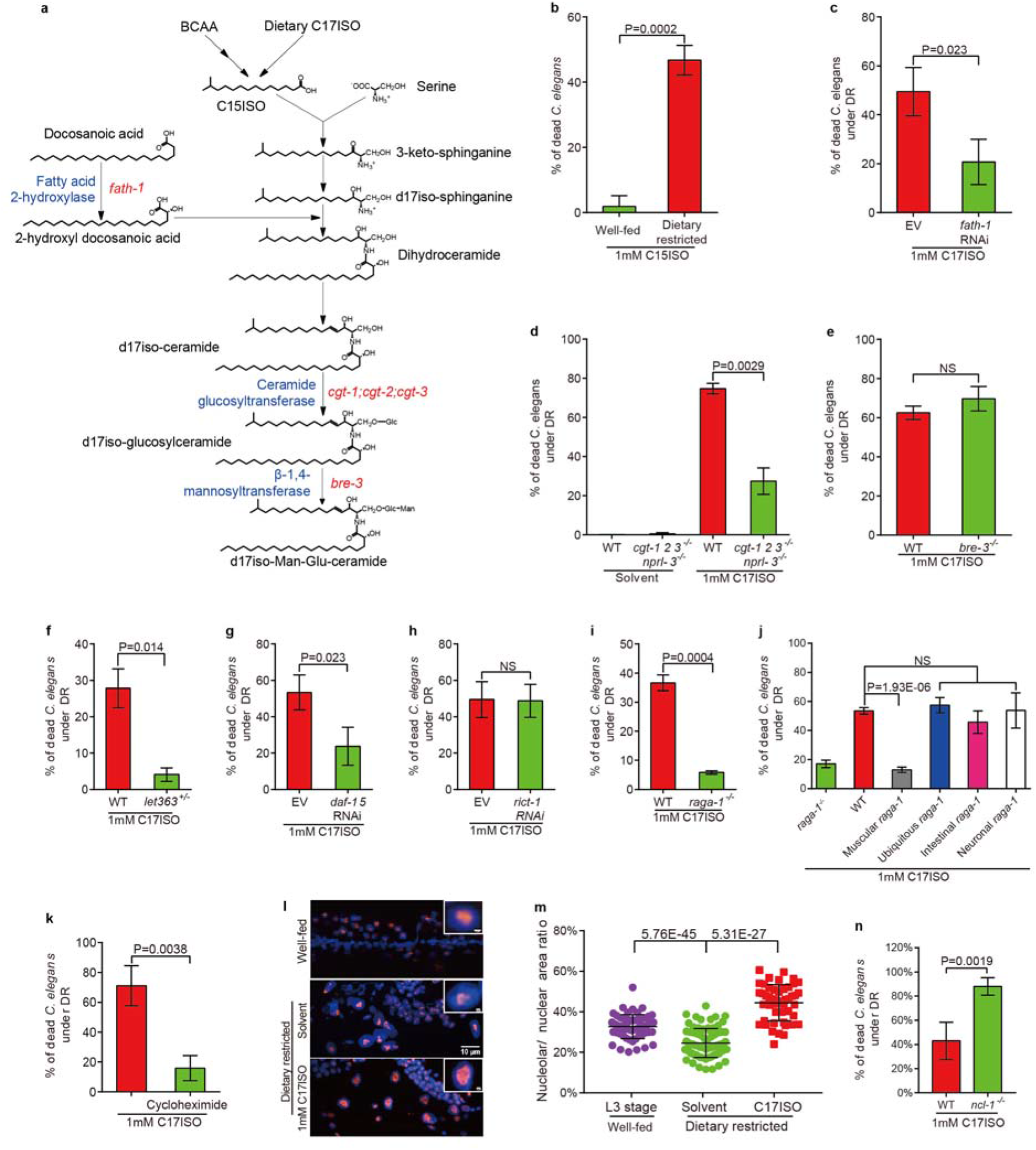
Glucosylceramide, mTORC1 and ribosomal biogenesis are critically involved in mmBCFA-mediated amino acid sensing. **a**, Cartoon illustration of a simplified, mmBCFA-involved sphingolipid biosynthetic pathway in *C. elegans.* The relevant enzymes and coding genes were labeled in blue and red, respectively. **b**, C15ISO, which can be derived from C17ISO, also induced the DUPED phenotype. **c-e**, Bar graphs showing that C17ISO induced DUPED was suppressed by RNAi of *fath-1* (disrupt dihydroceramide biosynthesis) or mutating all 3 genes for ceramide glucosyltransferases *(cgt-1/2/3(-);nprl-3(-)* quadruple mutant) (block glucosylceramide biosynthesis), but not by mutating *bre-3* (block mannosylglucosylceramide biosynthesis). f-i, Bar graphs showing that C17ISO-induced DUPED is suppressed in a *let-363/mTOR(-/+)* heterozygous mutant, RNAi knockdown of either *daf-15/raptor,* or a *raga-1/RagA/B(-)* mutation, but not by RNAi knockdown of *rict-1/RICTOR.* **j**, The dramatic suppression of C17ISO-induced DUPED by the *raga-1(-)* mutation was reversed (restored the high death rate) by transgenic expression of the wild type *raga-1* gene behind a ubiquitous, intestine-specific, or neuron-specific promoter, but not behind a muscle specific promoter. **k**, Protein synthesis inhibitor cycloheximide effectively suppressed C17ISO-induced DUPED. **l-m**, Representative fluorescent microscopic images and statistic data showing that the relative nucleolar size was significantly reduced under the DR condition, but was prominently increased with C17ISO supplement. n, Bar graph showing that an *ncl-1(-)* mutation significantly enhanced mmBCFA induced DUPED.

We then asked which signaling pathway could be responsible for the C17ISO/GlcCer-mediated AA sensing and developmental regulation in *C. elegans*. Although the Insulin/insulin-like growth factor signaling (IIS) and AMP-activated protein kinase (AMPK) pathway were known to be involved in the developmental growth/diapause decision making process under various DR conditions^13,19,31^, our tests using the C17ISO induced DUPED condition suggest that mutations in these two pathways *(age-1/PI3K, daf-16/FOXO* or *aak-2/AMPK)* do not play a major role in C17ISO/GlcCer-mediated AA sensing (Extended Data Fig. 3b-d). We then tested the role of mTOR signaling, a conserved pathway which acts as the central hub of amino acid sensing among eukaryotes^32^. In many organisms including *C. elegans,* partial decrease of mTOR pathway is critically involved in DR-induced lifespan extension^33,34^, suggesting mTOR activity was regulated by the nutrient condition. Moreover, we previously found that endogenous mmBCFAs and derived GlcCer are essential for maintaining mTORC1 activity in promoting *C. elegans* early larval development and foraging behavior^23,26^. Therefore, it is plausible that mTOR is the key signaling system that senses amino acid deprivation under the DR condition leading to protective arrest, and may sense C17ISO and promote downstream activities that resulted in the DUPED phenotype under DR. In this case, the DUPED phenotype would be expected to be suppressed by reducing mTOR activity. Because the *let-363/TOR* null mutant is lethal, we used a *let-363(-/+)* heterozygous mutant to lower mTOR activity^34^ Strikingly, the DUPED phenotype was dramatically suppressed in the mutant, suggesting that dietary C17ISO-induced lethality depends on high mTOR activity (Fig. 3f). mTOR can form two complexes, mTORC1 and mTORC2, and mTORC1 is mainly responsible for mediating amino acid sensing^18,35–37^. *daf-15/Raptor* and *rict-1/Rictor* encode key components of mTORC1 and mTORC2, respectively^18^. RNAi treatment of *daf-15/Raptor,* but not *rict-1/Rictor,* suppressed C17ISO-induced DUPED (Fig. 3g,h). Moreover, a loss-of-function mutation of *raga-1* /RagA/B, which was shown to play a critical role in mTORC1-dependent amino acid sensing in the cultured cells^26,38^], also strongly suppressed C17ISO-induced DUPED (Fig. 3i). These data indicate that the RagA/B-mTORC1 pathway plays a critical role in C17ISO-induced DUPED and is the key signaling system that triggers the protective arrest in response to amino acid shortage under the DR condition.

To determine the tissue where mTORC1 activity is sufficient for C17ISO-induced DUPED, we overexpressed the WT *raga-1* under various tissue-specific promoters in *raga-1(-)* and found intestinal or neuronal-specific expression of *raga-1* could restore the DUPED phenotype (Fig. 3j). Restoration of *raga-1* in other tissues such as muscle did not have a similar effect (Fig. 3j). These data suggest activating mTORC1 in either the intestine or neurons is sufficient to disrupt the protective function under the DR condition, which is different from its previously identified role in promoting L1 development [only intestinal mTORC1 is required^26^]. Assuming that the RAGA-1 protein acts cell-autonomously to activate mTORC1, these data also suggest that holding a low mTORC1 activity in both the intestine and neurons is required for the protective arrest of *C. elegans* larvae under the DR condition. Therefore, the mmBCFA-mediated amino acid sensing appears to involve the coordination of nutrient sensing by mTORC1 in the tissue where dietary nutrients are first received and the tissue where the regulatory instruction may be passed to other tissues.

Because we found supplementation with the AA mix could strongly suppress C17ISO induced lethality (Fig. 2b), we hypothesized that the lethality was caused by amino acid depletion from a cellular process downstream of mTORC1. One of the well-established mTORC1 functions is to suppress lysosomal autophagy, an important process for animals to recycle useless organelles and proteins to regenerate AA under DR condition^39^. However, we found that C17ISO supplement did not significantly affect the number of autophagosomes marked by LGG-1:: GFP (*C. elegans* homolog of ATG8)^40^ under the DR condition (Extended Data Fig. 3e,f). Moreover, the nuclear level of transcription factor PHA-4 (*C. elegans* FOXA), which is required for DR-induced autophagic response^41,42^, was also unchanged under C17ISO supplement (Extended Data Fig. 3g,h). These data indicate that C17ISO-induced DUPED was not mainly due to the inhibition of autophagy by mTORC1 under the DR condition.

Another well-established function of mTORC1 is to promote *de novo* protein biosynthesis^2^. We thus used cycloheximide (a translational elongation inhibitor to block protein synthesis)^43^ and found it dramatically suppressed the C17ISO induced DUPED phenotype (Fig. 3k). These data strongly suggest that C17ISO-mTORC1 signaling promoted unhealthy protein synthesis under the DR condition, which likely exhausted the limited amino acid pool.

mTORC1 could promote protein synthesis via ribosome biogenesis, which occurs in the nucleolus^44,45^. In eukaryotes including *C. elegans*, a cellular hallmark of ribosome biogenesis (such as Pol I-mediated rDNA transcription) is the nucleolus size^46^. Previous work in *C. elegans* showed *eat-2* induced DR or mTORC1 inactivation result in reduction in nucleolar size^47^. Indeed, we found in our DR animals that the average nucleolar size (marked by *C. elegans* nucleolar protein fibrillarin FIB-1) was significantly decreased (Fig. 3l,m). Strikingly, C17ISO supplementation almost doubled the nucleolar size of worms under DR (Fig. 3l,m). Moreover, *ncl-1* encodes a translational suppressor of fibrillarin of which a loss-of-function mutation is known to enlarge the nucleolar size and promote ribosomal genesis^46^. We found an *ncl-1(-)* mutation also significantly augmented the lethality of mmBCFA induced DUPED (Fig. 3n). These data indicate that high-level ribosome biogenesis is a critical cause of C17ISO-induced DUPED, further supporting that mmBCFA mediates the amino acid sensing by mTORC1 pathway.

We next explored if these observations were conserved in mammalian cultured cells. Like *C. elegans*, humans can both synthesize mmBCFAs from branched-chain AA through endogenous pathways and ingest dietary mmBCFAs through dairy product and meat^48,49^. In addition, similar to *C. elegans*, human gut bacteria metabolize BCAA and produce various short chain mmBCFA^50^. These suggest mmBCFA may also activate mTOR in mammals. To replicate the DR condition, we treated serum-deprived 3T3L1 cells with C17ISO and found it significantly enhanced the mTOR activity measured by P70S6K phosphorylation (Fig. 4a). In fact, when supplied at the same concentration, C17ISO activated mTOR even stronger than leucine, and such activation was completely suppressed by the mTORC1 inhibitor rapamycin (Fig. 4b). Moreover, GlcCer biosynthesis inhibitor AMP-Deoxynojirimycin (AMP) also significantly blocked the leucine and C17ISO induced P70S6K phosphorylation (Fig. 4b). These data not only suggest that mmBCFA also activates mTORC1 in mammalian 3T3L1 cells, but also suggest mmBCFA or even leucine induced mTORC1 activation partly depends on GlcCer biosynthesis, at least in certain mammalian cells. Taken together, we propose a model describing that *C. elegans* use mmBCFA as a major intermediate signal of the amino acid level to regulate mTORC1-dependent ribosomal genesis, protein biosynthesis and development, and a similar pathway may also exist in mammals (Fig. 4c).

**Fig 4.**
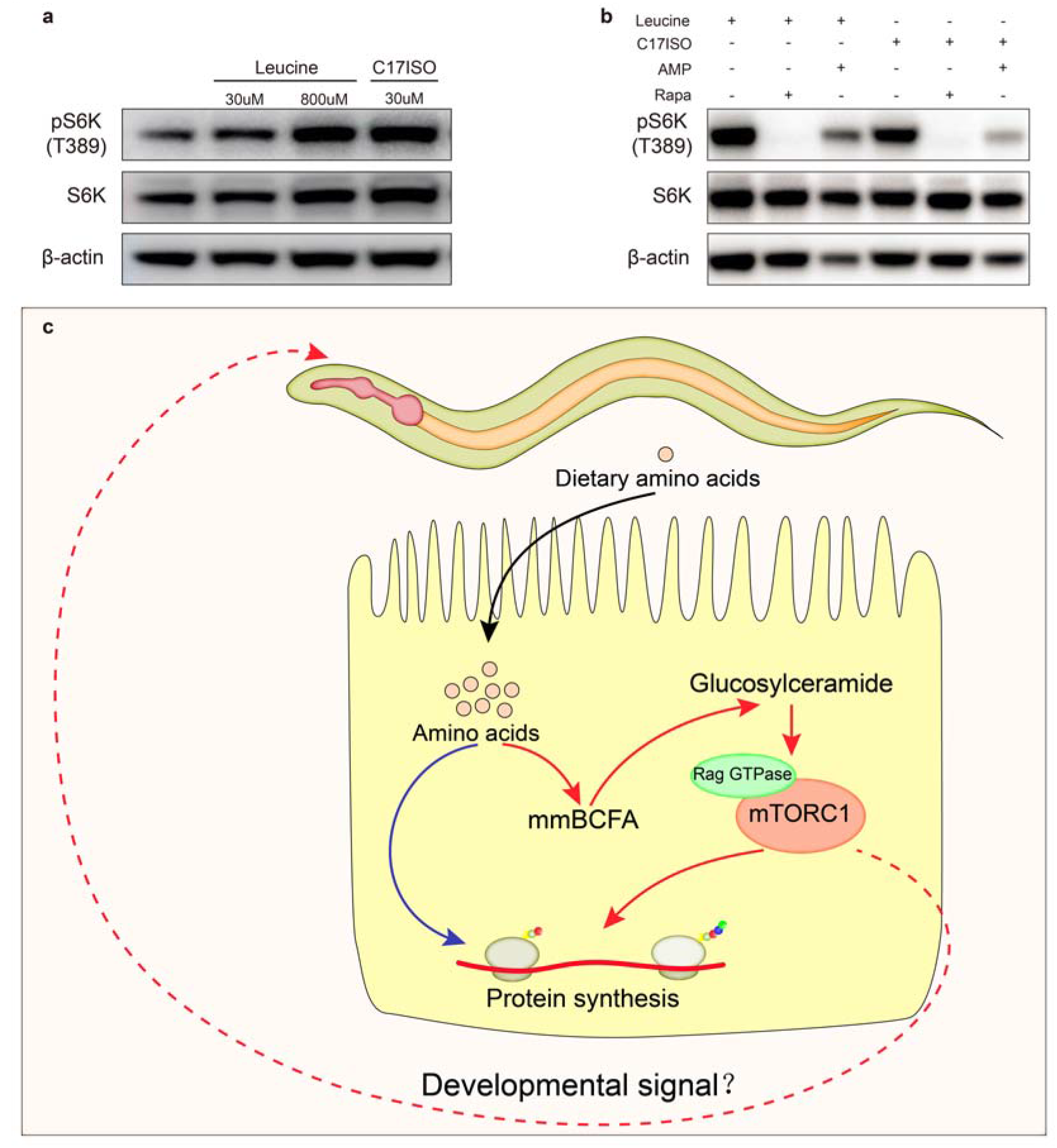
mmBCFA mediates the AA-mTORC1 pathway in mammalian 3T3L1 cells. **a-b,** Western blot image showing levels of S6K and phosphorylated S6K in cultured 3T3L1 cells under indicated condition. (**a**) C17ISO induced strong S6K phosphorylation under the serum starvation. (**b**) Ceramide glucosyltransferase inhibitor AMP-Deoxynojirimycin partly suppressed the leucine or C17ISO induced S6K phosphorylation under the serum starvation. **c,** A model showing dietary mmBCFA, which are derived from branched-chained amino acids and precursor of mmBCFA-containing glucosylceramide, critically mediates the sensing of overall AA level by mTORC1 in regulating *C. elegans* protein synthesis and development.

The mmBCFA-mediated AA sensing pathway we discovered in *C. elegans* shares many similarities with the mammalian counterparts in cultured cells. First, the mmBCFA signal to mTORC1 that results in DR lethality is dependent on RAGA-1 (*C. elegans* RagA/B) and DAF-15 (*C. elegans* RAPTOR), and both are key components in the mammalian amino acid sensing process^1^. Second, mmBCFA C17ISO is a metabolite directly derived from leucine, one of the most important essential amino acids in the mTORC1-dependent amino acid sensing pathway in mammals^2^. Last, the essential amino acid mixture which we found further enhanced the mmBCFA induced DUPED lethality, contains leucine, arginine and methionine that partially activate mTORC1 in mammals^5–7^. Given the broad existence of mmBCFA in human food/gut bacteria metabolites and the conserved mmBCFA/sphingolipid pathway in humans^21,48,51^, and our finding that mmBCFA C17ISO also activates mTORC1 in mammalian cells (Fig. 4C), further studies of this mmBCFA-mTORC1 regulatory function in humans and the potential therapeutic effect on mTORC1-related diseases should be explored. In addition, our model does not exclude the involvement of other nutrient sensing pathways in regulating development under the DR condition. In fact, the incomplete lethality caused by mmBCFA supplementation and incomplete developmental restoration by AA supplementation suggest that other nutrients and sensing pathways may also exist to regulate animal development.

## Acknowledgments

We thank Shohei Mitani, Yidong Shen, Shiqing Cai, Di Chen, Hongyun Tang, Quanjiang Ji, Szecheng J. Lo and the *C. elegans* Knockout Consortium and CGC (funded by NIH [P40OD010440]) for strains and advice. We thank the core facilities of ShanghaiTech School of Physical science and technology and School of Life science and technology for assistance; and our laboratory members for helpful discussions.

## Author contributions

MZ, FT Conception and design, Acquisition of data, Analysis and interpretation of data, Drafting or revising the article; NL, LZ Acquisition of data, Analysis and interpretation of data, Drafting or revising the article; Contributed unpublished essential data or reagents; HZ, Supervised the study, Conception and design, Drafting or revising the article.

## Data and materials availability

All data is available in the main text or the supplementary materials.

