## Supplemental figures and methods for "Monomethyl branched-chain fatty acid mediates amino acid sensing by mTORC1"

Materials and Methods

***Caenorhabditis elegans* strains and maintenance**

The following strains were obtained from the *Caenorhabditis* Genetics Center (CGC) or as indicated; wild type N2 Bristol, *raga-1(ok386)*, *glp-1(e2141)*, *glp-1(ar202)*, *ife-2(ok306)*, *bre-3(ye26)*, *rsks-1(ok1255)*, *let-363(ok3018)/hT2[bli-4(e937)let-2(q782)qIs48]*. The *cgt-1(tm1027)*, *cgt-2(tm1192)* and *cgt-3(tm504)* mutants were provided by the Mitani Lab (National BioResource Project, Tokyo, Japan). *C. elegans* were maintained at 20°C on NGM plates (referred to as standard plates) with *E. coli* OP50 as the bacterial food (OP50/NGM). The bleaching process was modified from the standard protocol by two more M9 washes ^1^.

**Dietary macronutrient preparation**

Fatty acids C15ISO, C17ISO (Larodan), C18:1n7, C20:5n3 (Cayman), palmitic acid (Sigma) and stearic acid (TCI) were dissolved in DMSO and prepared as 10mM stocks. Leucine, valine, lysine, isoleucine, histidine, arginine, tyrosine (Sigma), glucose, sucrose, fructose (Sangon), were prepared as 100mM stocks in water. A stock solution was diluted 10 times with OP50 bacterial suspension. The recipe for amino acid mixture (Sigma) was the same as a previously reported chemically defined *C. elegans* medium (CeMM)^2^. For EAA, NEAA and AA mix supplemented condition, 850μl of 1x EAA mixture solution, 1.7ml of 1x NEAA mixture or 850μl EAA plus 1.7ml NEAA mixture was added on the carbenicillin containing NGM plate respectively, and dried before seeding *E. coli.*

**Preparation of dietary restriction and concentrated food plates**

A single colony of OP50 bacteria was inoculated into ~70ml liquid LB broth and cultured at 37°C overnight. The bacterial culture was then diluted with fresh LB broth to the concentration of OD600=0.35 (about 5.8 x 10^8^cfu/ml). 350μl of such bacterial culture was seeded on a carbenicillin NGM plate (with 50ng/ml carbenicillin, named antibiotic plate hereafter) and this diluted food (2 x 10^8^ cfu) was used as the dietary restriction feeding condition. For the “well-fed” plates (Figure 1L), 350μl of 10-fold concentrated bacterial culture was seeded on the same antibiotic NGM plate (2 x 10^9^ cfu) and was considered well-fed condition.

***C. elegans* reversible arrest on the dietary restricted plate**

50~70 synchronized L1 worms were added to the dietary restricted plates and cultivate in 20°C for 3 days. The developmental stage of each worm was scored. To verify whether such developmental arrest was reversible, at day 6, those L3 arrest worms were transferred back to standard OP50 seeded NGM plates and their developmental stages were scored 2 days later.

**Macronutrient signal screening assay**

Macronutrient supplement was mixed with the diluted OP50 bacteria (OD600=0.35) (described above) to a final volume of 350μl before they were seeded on the antibiotic NGM plates. 50~70 synchronized L1 worms were added to those plates and cultivated at 20°C for 6 days, then the survival ratio and physiological defects were scored.

**C17ISO-induced death under poor environmental diet (DUPED)**

This assay is essentially the same as the one for macronutrient signal screening. We slightly modified the assay under various experimental conditions (described below).

C15ISO-induced lethality under dietary restriction: The well-fed plates contained 350μl (1.9 x 10^10^cfu/ml) bacteria, and the lethality ratio was scored at day 4 instead of day 6, to prevent bagging-induced lethality.

For assays using additional nutrients to suppress C17ISO-induced DUPED, 100μl 10mM palmitic acid, 100μl 100mM glucose, or 2.55ml amino acid mixture (Lu and Goetsch 1993) were added to the antibiotic NGM plates.

For assays using worms started at various developmental stages (described in Sup Fig. 1A): Synchronized L1 worms were cultivated on standard NGM plates at 20°C. Then those worms were transferred to DR plates with 1mM C17ISO [25uM 5-fluro-2’-deoxyuridine (FUDR) added in the NGM plates, if indicated] at the indicated developmental stages (L1, L3 or day 4 adult stage). At day 4, live worms were transferred to a new plate to prevent bagging. The survival ratio (survivors/ [all worms on the plate plus dead worms before the plate transfer]) was scored on day 6.

For the DUPED assays using *glp-1* temperature sensitive mutants, day 1 adult worms (6-10 worms/plate) with indicated genotypes [*glp-1(e2141) or glp-1(ar202)*] were grown at 15°C on standard NGM plates and their adult progeny were bleached and synchronized in M9 at 25°C for 20 hours. Those L1 worms were grown on indicated plates at 25°C instead of 20°C.

**Lifespan assay**

For the lifespan assay under the well-fed condition described in Figure 1M, we used a previously reported method ^3^: Briefly, synchronized L1 worms were cultivated on standard NGM plates at 20°C for about 3 days. Then day 1 adults were transferred to plates (30 worms per plate) containing 1mM C17ISO or DMSO solvent control. Worms were judged as dead when they did not respond to repeated prodding with a pick and had no pharyngeal pumping. Dead worms were counted daily. Worms that crawled off plates were not counted.

For the lifespan assay under the dietary restriction condition (described in Fig. 1K): the method was modified so that synchronized L1 stage worms were grown on dietary restricted plates (30 worms per plate) and were transferred to a new dietary restricted plate every 3 days to prevent the depletion of food.

For the lifespan assay under the dietary restriction condition from the adult stage (described in Supp. Fig 1C): the assay was modified from a previously described one ^4^. Briefly, diluted bacterial culture (5 x 10^8^ cfu/ml) was mixed with 10mM C17ISO (dissolved in DMSO) or solvent only at 10:1 ratio. 166μl of this mixture solution was seeded on the NGM plate (60mm) containing 50ng/ml carbenicillin. The lifespan assay of worms grown on standard OP50 NGM plates was the same as previously described ^3^.

**RNAi feeding assay** （under DR condition）

All *E. coli* strains for feeding RNAi were from our previous work ^5^*(daf-15)* or the ORF-RNAi library (Dharmacon, Horizon discovery) *(fath-1* and *rict-1)*. The strains were confirmed by sequencing. Preparation of RNAi NGM plates was done according to the standard protocol ^6^. Briefly, day 1 adult (P0) worms were transferred to the indicated RNAi plates (8/plate). F1 adults were collected, bleached and synchronized. Those synchronized L1 worms (F2) were transferred to DR +1mM C17ISO plates (without RNAi bacteria). Survival was determined at day 3 *(daf-15)*, day 4 *(fath-1)* instead of day 6 to prevent depletion of RNAi effect. Worms that crawled off the NGM plates were excluded.

***raga-1(ok386)* rescue experiment**

Whole genomic *raga-1* ORF DNA sequence was cloned and driven by the following promoters: *Pges-1*(intestine-specific), *Prgef-1*(neuron-specific), *Pmyo-3*(muscle-specific), *Prpl-28*(ubiquitous), and injected into the *raga-1(ok386)* mutant with the *Pmyo-2::gfp* co-injection marker. Animals carrying the marker were considered transgenic strains.

**Autophagy evaluation**

The autophagy level was evaluated by scoring the puncta numbers of the LGG-1::GFP translational reporter^7^. Synchronized L1 worms were transferred to DR + 1mM C17ISO plates (or +DMSO as the solvent control) and cultivated at 20°C. After 3 days, the number of LGG-1::GFP positive puncta in epidermal seam cells were counted in L3 worms. For each condition, 538 seam cells (64 worms) were analyzed.

**Western blot assay for P70S6K phosphorylation in 3T3-L1 cells**

This method was adopted from a published report ^8^. Briefly, 3T3-L1 cells were cultured in DMEM supplied with 10% new calf serum (NBS) (37ºC, 5% CO_2_) for 2 days, and then transferred to serum and BCAA-free DMEM (BasalMedia) for 2 h. This was followed by the addition of various nutrients [Leucine (800μM in Fig. 4B], C17ISO [30μM in Fig. 4C] and/or mTORC1 inhibitor Rapamycin [final concentration 0.1μM] into the medium. After 2 hours, the cells were harvested and protein was extracted for Western-Blot assay. The total level of P70S6K and phospho-P70S6K were measured by using relevant antibodies (Cell Signaling #9202 and #9234).

In the GlcCer biosynthesis inhibition experiment, GlcCer synthase inhibitor AMP-Deoxynojirimycin was added into the NBS/DMEM medium (final concentration 10uM] 48 hours before the cells were shifted to the Serum-free DMEM (also contain 10 uM AMP-Deoxynojirimycin).

**Immunofluorescence**

The immunofluorescence assay of FIB-1 was adopted from a previously published “frozen crack” method ^9^. Briefly, L3/L4 animals were collected and washed three times with distilled water to remove bacteria. The worms were then transferred and spread to the center of poly-lysine coated slides and a cover slide was placed on top. Appropriate pressure was applied to ensure that the worms contacted with both slides and this pair of slides were transferred to the top of a piece of metal on dry ice for 30min. Then the slide pair was swiftly twisted apart, and the polylysine coated slide was immediately subjected to fixation (4% Paraformaldehyde [PFA] for 30min at room temperature). After the fixation, the samples were washed three times using PBS and then blocked with 0.5% Normal Donkey serum in PBS with 0.1% Triton-100 for 1 hour at room temperature. The samples were then washed three times in PBS with 0.1% Tween20 (PBST). After blocking, the samples were incubated with a primary antibody against Fibrillarin (Abcam ab4566,1:400) overnight at 4℃. After three washes in PBST, the samples were incubated with a secondary antibody (Invitrogen A11005, 1:800) for 1 hour at room temperature followed by three washes in PBST, then stained with DAPI for 5 min. The samples were mounted with Antifade Mounting Medium (Beyotime P0126). 126 gut cells (36 animals) were analyzed.

**Microscopy**

Autophagy analysis (LGG-1::GFP) were completed using Nikon CSU SORA spinning disk microscope and a prime 95B photometrics sCMOS camera. Plate phenotypes were observed with a MVX-ZB10 fluorescence dissecting microscope and a BioHD-C20 CMOS camera. Nucleolar area was quantified with Fiji software.

**Statistical analysis**

All statistical analysis, except lifespan related experiments, were preformed using Student’s *t*-test and p<0.05 was considered a significant difference. For lifespan-related assays, log-rank (mantel-Cox) tests were used to calculate the *p* values and p<0.05 was considered a significant difference.

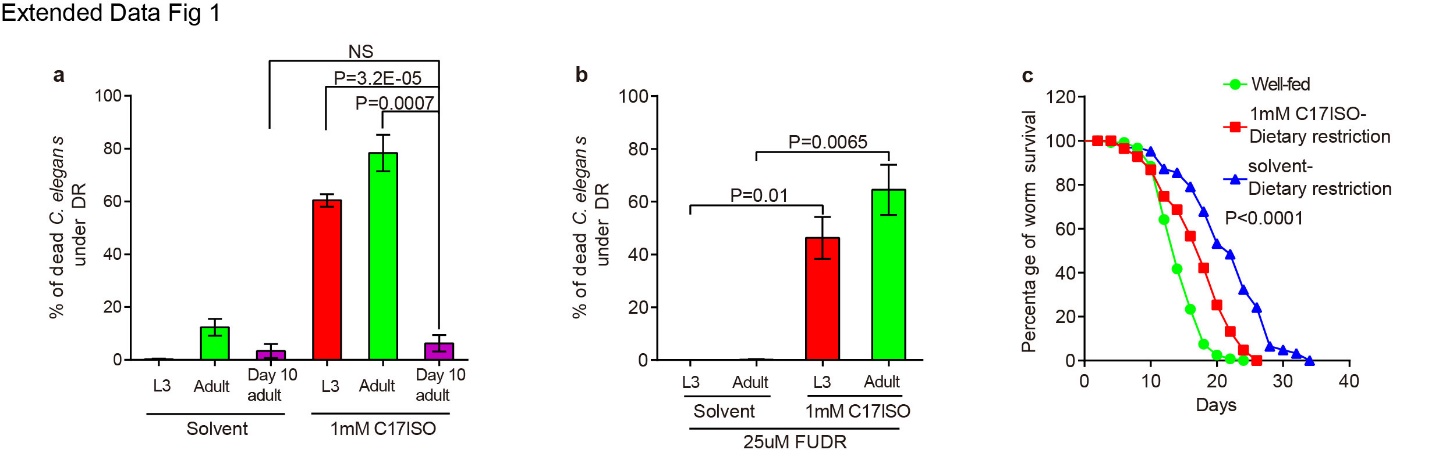

Extend Data Fig. 1| Summary of C17ISO effect on different stage *C. elegans*. a-b, Bar graphs showing the percentage of death in *C. elegans* under indicated feeding conditions at different developmental stages. a, C17ISO induced the strong DUPED phenotype in L3 and young adult animals, but did not have an obvious effect on the older adult animals. b, The DUPED phenotype was not suppressed by the addition of FUDR (germline proliferation inhibitor), suggesting that the lethality was not due to inability of the worms to lay eggs. c, Survival curves showing *C. elegans* life span under the DR +/- C17ISO (supplemented since day 4 adults). C17ISO supplement moderately but statistically significantly shortened the life span of *C. elegans*.

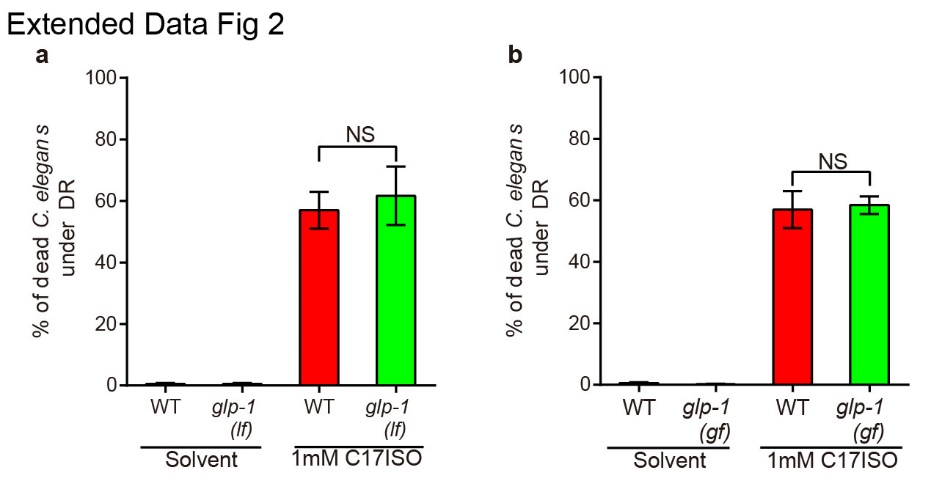

Extend Data Fig. 2 | DUPED phenotype was independent of gonad. a-b, Bar graphs showing the percentage of death of *C. elegans* strains under indicated feeding conditions. a, The Notch receptor GLP-1 protein is the master regulator of germline mitotic proliferation ^10^. The C17ISO-induced DUPED phenotype is not suppressed by blocking germline proliferation with a *glp-1* loss-of-function mutation [*glp-1(e2141)* or *glp-1(-)*]. b, The phenotype is also not enhanced by hyper-activation of germline proliferation in a *glp-1* gain-of-function allele [*glp-1* *(ar202),* or *glp-1(gf)*]*.*

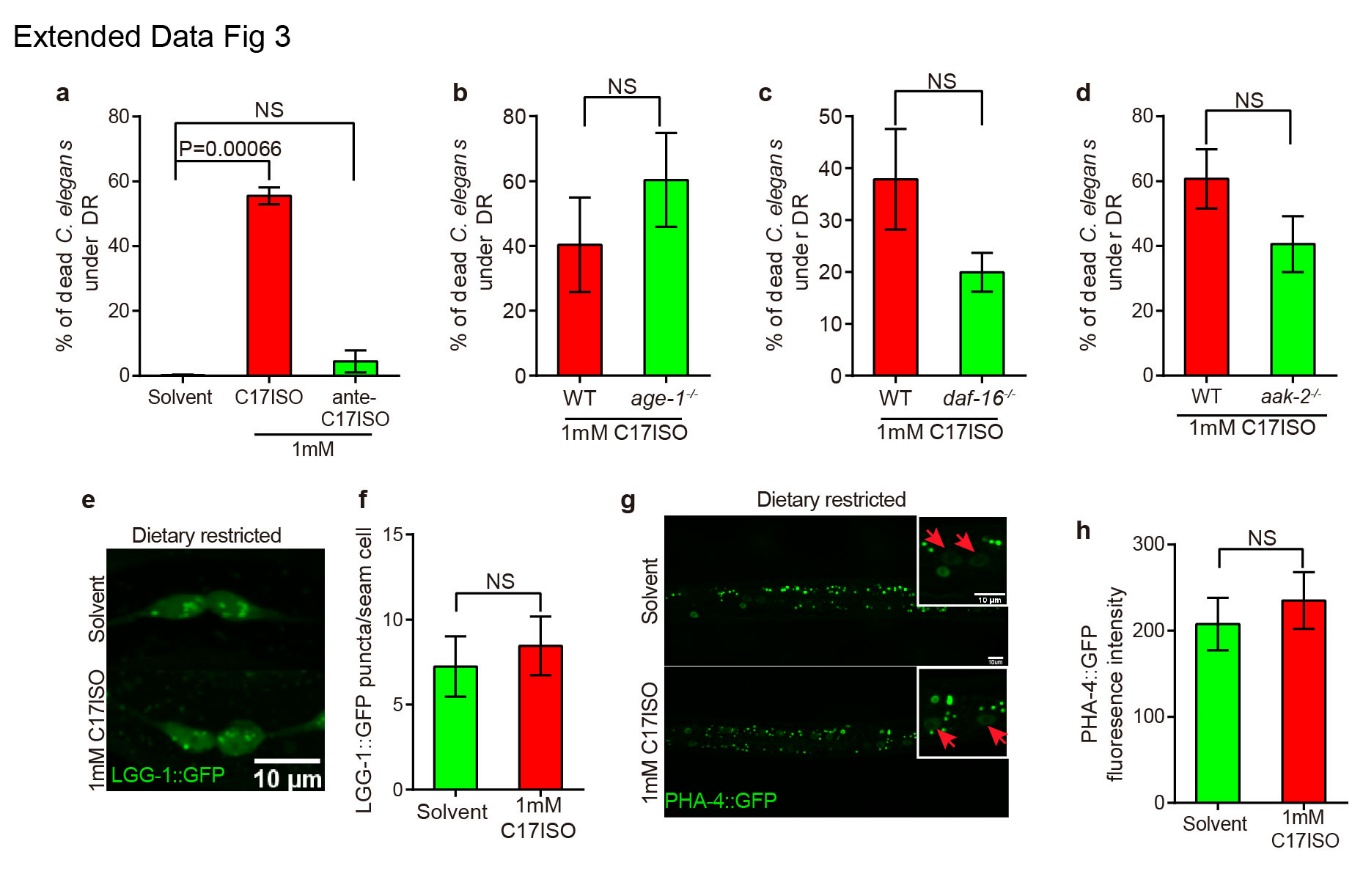
Extended Data Fig. 3 | Summary of C17ISO effect on canonical developmental pathway. a-d, Bar graphs showing the percentage of death of various *C. elegans* strains under indicated feeding conditions. a, C17anteISO did not trigger the DUPED phenotype. The C17ISO-induced DUPED phenotype was not strongly suppressed by loss of *age-1* (b), *daf-16* (c), or *aak-2* (d). e-f, Representative fluorescent microscopy images and statistical data showing that the number of autophagosome puncta marked by LGG-1::GFP in WT seam cell was not significantly altered by C17ISO supplementation. g-h, Representative fluorescent microscopy images and statistical data showing nucleic PHA-4 level marked by Prom*_pha-4_*-PHA-4::GFP in WT intestine cells (marked by arrows) was not altered by C17ISO supplementation.

­
